# Neural Correlates of Multisensory Reliability and Perceptual Weights Emerge at Early Latencies during Audio-visual Integration

**DOI:** 10.1101/116392

**Authors:** Stephanie C. Boyle, Stephanie J. Kayser, Christoph Kayser

**Author notes:** Corresponding author: C. Kayser. Institute of Neuroscience and Psychology, University of Glasgow, Hillhead Street 58, Glasgow, G12 8QB, UK. Email:_.

## Abstract

To make accurate perceptual estimates observers must take the reliability of sensory information into account. Despite many behavioural studies showing that subjects weight individual sensory cues in proportion to their reliabilities, it is still unclear when during a trial neuronal responses are modulated by the reliability of sensory information, or when they reflect the perceptual weights attributed to each sensory input during decision making. We investigated these questions using a combination of psychophysics, EEG based neuroimaging and single-trial decoding. Our results show that the weighted integration of sensory information in the brain is a dynamic process; effects of sensory reliability on task-relevant EEG components were evident around 84ms after stimulus onset, while neural correlates of perceptual weights emerged around 120ms after stimulus onset. These neural processes also had different underlying topographies, arising from areas consistent with sensory and parietal regions. Together these results reveal the temporal dynamics of perceptual and neural audio-visual integration and support the notion of temporally early and functionally specific multisensory processes in the brain.

## Introduction

The reliability of the information received by our senses varies. For example, visual cues become unreliable in dim or fogged conditions, and auditory cues become unreliable in loud or noisy situations. Studies have shown that observers deal with such variations in reliability by combining cues, where each is weighted in proportion to its apparent reliability [Battaglia et al., 2003; Butler et al., 2010; Ernst and Banks, 2002; Fetsch et al., 2009; Helbig and Ernst, 2007; Hillis et al., 2004; Jacobs, 1999; Raposo et al., 2012; Sheppard et al., 2013]. By doing so, more reliable cues are assigned a higher weight and have stronger influence on the perceptual estimate. In most cases, this leads to a more precise and reliable percept [Angelaki et al., 2009; Ernst, 2006; Ernst and Bülthoff, 2004; Fetsch et al., 2013; Rohde et al., 2015].

Despite many psychophysical studies investigating the weighted combination of sensory information, the neural mechanisms underlying this process remain unclear. Single-cell recordings have shown that neuronal sensory weights extracted from selected brain regions can vary with cue reliability in a manner consistent with predictions from statistical optimality [Gu et al., 2008; Morgan et al., 2008], and can predict those derived from behaviour [Fetsch et al., 2012]. Similarly, fMRI studies have demonstrated that BOLD responses are modulated by sensory reliability during visual-tactile [Beauchamp et al., 2010; Helbig et al., 2012] and audio-visual tasks [Rohe and Noppeney, 2016], and have suggested that the sensory weighting process emerges gradually along the cortical hierarchy [Rohe and Noppeney, 2015a; Rohe and Noppeney, 2016].

While providing valuable computational insights, these studies have not determined the temporal evolution of the neural processes implementing the weighting of sensory information. Studies comparing neural response amplitudes underlying sensory integration have shown that multisensory interactions can occur at surprisingly short latencies, starting around 40-76ms after stimulus onset [Cappe et al., 2010; Cappe et al., 2012; Fort et al., 2002; Foxe et al., 2000; Giard and Peronnet, 1999; De Meo et al., 2015; Molholm et al., 2002; Murray et al., 2004; Murray et al., 2016]. However these results were obtained by comparing generic response amplitudes between uni-and multi-sensory conditions, and hence did not specifically associate neural activity with either sensory reliability or a specific computational process during cue integration. Therefore it remains unclear when following stimulus onset neuronal responses are modulated by the reliability of sensory information, and when they reflect the sensory weights that drive the subsequent perceptual decision [Bizley et al., 2016].

In this study we investigated these questions by examining the temporal dynamics of weighted cue combination during an audio-visual task. We combined a rate discrimination task with EEG based neuroimaging, single-trial decoding, and linear modelling to identify the neural correlates of audio-visual cue weighting. Our results show that neural activity is modulated by sensory reliability early in the trial, starting from 84ms after stimulus onset. Furthermore, we find that neural correlates of perceptual weights emerge shortly after stimulus onset (around 120ms) and well before a decision is made. Finally, these EEG correlates of sensory reliability and perceptual weights have topographies that are consistent with activations in early sensory cortical and parietal brain areas. Taken together, these results suggesting that reliability based cue weighting computations occur early during the audio-visual integration process.

## Materials and Methods

### Subjects

We obtained data from 20 right-handed subjects (13 females; mean age 26 years) after written informed consent. The sample size was set a priori to 20, based on sample sizes used in related previous EEG studies and general recommendations [Simmons et al., 2011]. All subjects reported normal or corrected to normal vision, normal hearing, and received £6 per hour for their participation. The study was approved by the local ethics committee (College of Science and Engineering, University of Glasgow) and conducted in accordance with the Declaration of Helsinki.

### Stimuli and Task

The task was an adapted version of a 2-alternative forced choice rate discrimination task [Raposo et al., 2012; Sheppard et al., 2013]. Subjects were presented with two streams (each lasting 900ms) of auditory, visual or audio-visual events and asked to decide which stream had a higher event rate (Fig. 1A). Visual events were noise squares (3x3cm, 2.1° of visual angle, flashed for 12ms each) presented atop a static pink-noise background image. Acoustic events were brief click sounds (65 dB SPL, 12ms duration) presented in silence. Individual acoustic or visual events were instantiated by random pauses of short (48ms) or long (96ms) intervals, causing them to appear as auditory and/or visual flicker.

**Figure 1.**
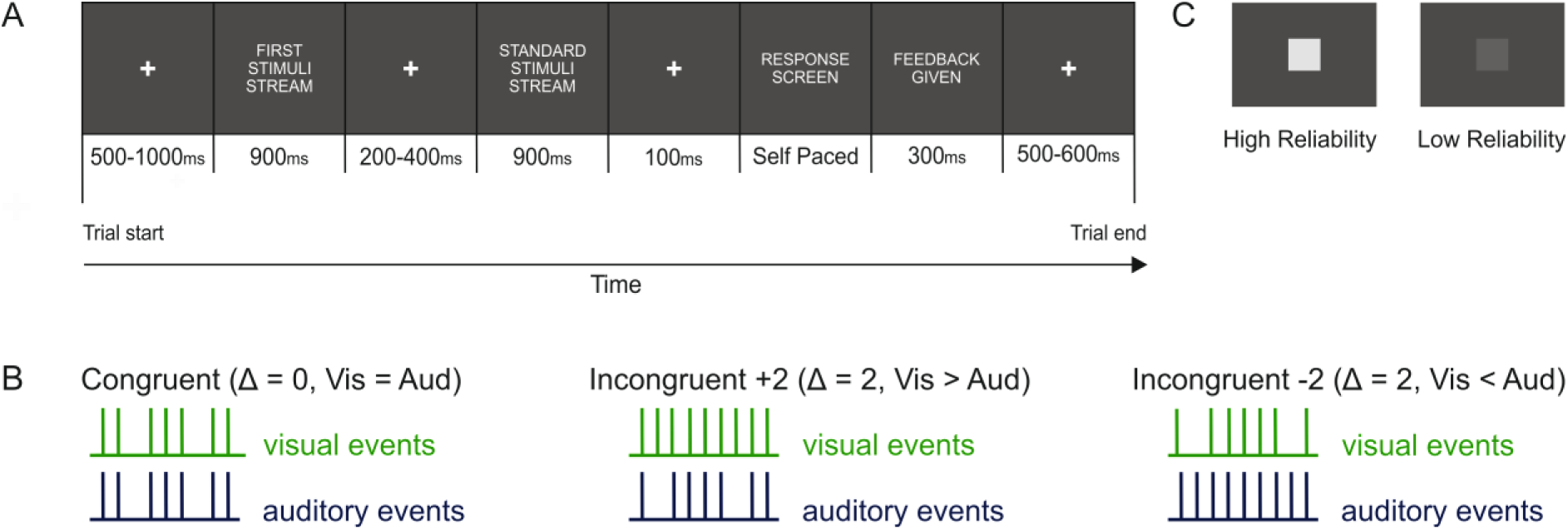
Experimental set up. (A) Subjects were presented with two sequential streams of auditory, visual and/or audio-visual events and had to indicate which stream contained more events. The first stream varied in modality, event rate, reliability and congruency of the events (see Methods). (B) Schematic showing one combination for each level of congruency (left: equal rates, middle: auditory fewer events, right: auditory more events). Δ = Visual – Auditory rate. (C) Example of high and low reliability visual stimuli.

In the first “experimental” stream, events were presented at seven different rates (8 to 14Hz). In the second “standard” stream events always flickered at 11Hz. We also manipulated the reliability of the visual stimulus, as well as the congruency between the rates of the auditory and visual stimuli [Angelaki et al., 2009; Fetsch et al., 2012; Fetsch et al., 2013; Sheppard et al., 2013]. Placing the audio-visual cues in conflict is necessary, as it allows assessment of the degree to which subjects are biased towards each cue.

Congruency was manipulated by introducing differences in the event rate between the auditory and visual streams. Audio-visual trials were either Congruent (Δ = 0) with auditory and visual streams each having the same number of events, or incongruent, with the visual either containing two more (Δ =+2), or two fewer (Δ = -2) events than the auditory stream (Fig. 1B).

The reliability of the visual stimulus was manipulated by adjusting the contrast of the visual stimulus relative to the background (Fig. 1C). Contrast levels were derived to match individual subject’s psychometric thresholds in separate calibration blocks carried out prior to the main experiment (see *Procedure).* Auditory reliability was constant throughout.

These manipulations resulted in: three unisensory conditions (auditory [AUD], visual high [VH] and visual low [VL]), two congruent (Δ =0) multisensory conditions (one where both the streams were highly reliable [AVH], and one where the auditory had high and the visual low reliability [AVL]) and four incongruent audiovisual conditions (AVH Δ =+2, AVH Δ =-2, AVL Δ =+2 and AVL Δ =-2).

### Experimental Procedure

The experiment was controlled through MATLAB (MathWorks) using the Psychophysics Toolbox Extensions [Brainard, 1997] and custom written scripts.

Auditory stimuli were presented using Sennheiser headphones and visual stimuli were presented on a Hansol 2100A CRT monitor at a refresh rate of 85 Hz. All recordings were carried out in a dark and electrical shielded room.

Subjects completed two simultaneous behavioural and EEG sessions (one session per day). Each session started with two unisensory calibration blocks used to calibrate performance between auditory and visual trials for each observer. The auditory calibration block consisted of 30 trials of auditory stimuli presented in silence (high reliable auditory stimuli), and an overall performance score was calculated. The visual calibration block consisted of 150 trials (30 trials x 5 SNRs), where the reliability of the visual stimulus varied systematically from high to low reliability. Psychometric functions were fit to the visual data, and two signal-to-noise (SNR) levels for visual reliability selected from the resulting psychometric curve. Visual high reliability was set as the SNR at which visual performance was equal to performance on the auditory calibration block. Visual low reliability was set at the SNR at which performance was ~30% lower than auditory performance.

Each block of the main study consisted of 510 trials with modality (auditory, visual, audio-visual), reliability (visual high and low), event rate (8-14Hz) and congruency (audio-visual Δ = 0, ± 2) varying pseudo-randomly across trials *(see Stimuli and Task).* In total, subjects completed four blocks across the two sessions, resulting in approximately 2040 trials per subject.

Each trial began with a white fixation cross presented centrally on a dark grey noise image (500-1000ms). This was followed by the first “experimental” stream (900ms), a fixation period (200-400ms), and then the standard stream (900ms). Subjects were then cued to respond using the left (“first stream has more events”) or right (“second stream has more events”) keyboard buttons, and received feedback on their performance (Fig. 1A). For trials where the rates in the experimental and standard stream were equal, feedback was randomly generated.

### EEG Recording and Preprocessing

EEG data was recorded using a 64-channel BioSemi system and ActiView recording software (Biosemi, Amsterdam, Netherlands). Signals were digitised at 512 Hz and band-pass filtered online between 0.16 and 100 Hz. Signals originating from ocular muscles were recorded from four additional electrodes placed below and at the outer canthi of each eye.

Data from individual subject blocks were preprocessed separately in MATLAB using the FieldTrip toolbox [Oostenveld et al., 2011] and custom written scripts. Epochs around the first stimuli stream (-1 to 2s relative to stream onset) were extracted and filtered between 0.5 and 90 Hz (Butterworth filter) and down-sampled to 200 Hz. Potential signal artefacts were removed using independent component analysis (ICA) as implemented in the FieldTrip toolbox [Oostenveld et al., 2011] and components related to typical eye blink activity or noisy electrode channels were removed. Horizontal, vertical and radial EOG signals were computed using established procedures [Hipp and Siegel, 2013; Keren et al., 2010] and trials during which there was a high correlation between eye movements and components in the EEG data were removed. Finally, remaining trials with amplitudes exceeding ±120 μV were removed. Successful cleaning was verified by visual inspection of single trials. For one subject (S20), three noisy channels (FT7, P9, TP8) were interpolated using the channel repair function as implemented in the FieldTrip toolbox.

## Analysis Methods

### Psychometric performance and Bayesian Integration Model

For each subject, modality, and stimulation rate, the proportion of “first stream had a higher event rate” responses were calculated and cumulative Gaussian functions fit to the data using the psignifit toolbox for Matlab [Fründ et al., 2011] http://psignifit.sourceforge.net/). The threshold (s.d., σ) and the point of subjective equality (PSE, μ) were obtained from the best fitting function (2000 simulations via bootstrapping), and used to calculate predicted and observed perceptual weights [Fetsch et al., 2012].

Predicted weights reflect the weights that a Bayesian optimal observer would assign to each sensory cue in multisensory conditions [Fetsch et al., 2012]. These were calculated using the thresholds (σ) from unisensory trials:

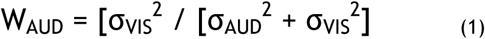

Observed perceptual weights represent the apparent weight a subject assigns to each sensory cue. These were calculated from the PSE (μ) from multisensory trials:

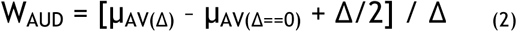
where Δ represents the incongruency between the auditory and visual stimuli [Fetsch et al., 2012]. For both perceptual and observed weights we assumed that auditory and visual weights sum to one:

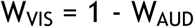

Predicted and observed weights were derived for each modality and reliability separately, and averaged over congruency levels.

### Time-dependent Perceptual Weights

To examine how perceptual weights evolved over the course of a trial, we used logistic regression to model the relationship between sensory evidence and behavioural reports at each time point.

As a measure of sensory evidence, we used the *accumulated rate,* defined as the number of stimulus events presented up to each time point in the trial. This was generated for each trial, condition (AVH auditory, AVH visual, AVL auditory and AVL visual) and time point separately, yielding a trial and time-specific measure of the experienced sensory evidence. This analysis was restricted to incongruent audio-visual trials and a time window from 24ms to 600ms post stimulus onset to account for null values (pre-24ms) and multicollinearity in the predictor matrix (post 600ms).

To assess whether the accumulated rate was a significant predictor of perceptual choice, we quantified the predictive performance of the regression model (referred to as Az) using the area under the receiver operator characteristic (ROC) of the regression model, based on 10-fold cross-validation (see *Statistics).* To determine how well the perceptual weights derived from the psychometric curves corresponded to the perceptual weights derived from the regression model, we also computed the correlation between the reliability influence for each pair of weights (psychometric and regression) at each time point during the trial (see *Statistics).* The reliability influence was here defined as the effect of visual reliability on auditory weights:

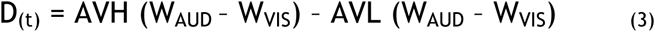

### Single-Trial EEG Analysis

We used single-trial, multivariate linear discriminant analysis [Kayser et al., 2016; Parra et al., 2005; Philiastides et al., 2006; Philiastides et al., 2014; Ratcliff et al., 2009] to uncover EEG components that best discriminated between our two conditions of interest. Here, we chose to discriminate between whether the first stream had an event rate that was lower or higher than the standard stream (</> 11 Hz), as this reflected the task the subjects were asked to complete. This analysis generated a one-dimensional projection (Y_t_) of the multidimensional EEG data (X_t_), defined by spatial weights (W_t_) and a constant (C):

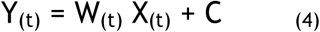

where the weight vector (W) represents the activity components most sensitive to the sensory stimuli, and the discriminant output (Y) provides a neural signature of the quality of the single-trial evidence about the condition of interest. This approach preserves the trial-to-trial variability of the data, and is assumed to be a better estimator of the underlying single trial task-relevant activity than the data on individual channels [Blankertz et al., 2011; Kayser et al., 2016; Parra et al., 2005; Philiastides et al., 2014].

Classification was based on regularised linear discriminant analysis [Philiastides et al., 2014], and applied to the EEG activity at each 5ms time point from stimulus onset to 600ms post stimulus onset, in sliding time windows of 55ms. For each time point, the discriminant output (Y) was aligned to the onset of the 55ms window. To avoid introducing bias to either sensory modality, we derived the weighting vector (W_t_) and constant (C) from the congruent audio-visual trials only (AVH and AVL Δ=0), and applied these to all other trials at the same time point. Scalp topographies for the discriminating component were estimated via the forward model [Philiastides et al., 2014], defined as the normalised correlation between the discriminant output and the EEG activity.

### Neural Weights

To quantify the apparent weight with which the sensory information in each modality contributed to the neural representation of the event rate we used linear regression. Similar to the behavioural data, the trial specific accumulated rates were used as predictors and regressed against the discriminant output (Y) at each time point (24ms to 600ms) in the trial. As conflict between the sensory cues is necessary to see how subjects are weighting the sensory information, this analysis was restricted to incongruent audio-visual conditions (AVH and AVL Δ ±2) and included separate weights for each modality in the high and low reliability conditions. This resulted in four neural weights for each time point (one for AVH auditory, AVH visual, AVL auditory and AVL visual). To assess the relationship between these neural weights and the time-dependent perceptual weights we correlated the reliability influence (Eqn. 3) between these at each time point.

### Statistics

All descriptive statistics reported represent median values. All Z values reported were generated from a two-sided Wilcoxon signed rank test after testing assumptions of normality, and effect sizes calculated by dividing the Z value by the square root of N (where N = the number of observations rather than subjects). Correlations were calculated using Spearman rank correlation analysis. All reported p-values were checked for inconsistencies using the R software package “statcheck” [Nuijten et al., 2015].

To correct for multiple comparisons along time, data were shuffled randomly across conditions, and for each separate comparison a distribution of t-values based on 1,000 randomisations was computed. Significance levels were determined using a cluster based randomisation technique [Maris and Oostenveld, 2007] using a cluster-threshold of t=1.8 and the max-size as cluster-forming variable (referred to as cluster randomisation test in text).

Significance levels of classification performance (Az) were determined by randomly shuffling data by condition 2000 times, computing the group averaged Az value for each randomisation, and taking the maximal Az value over time to correct for multiple comparisons (referred to as randomisation test in text). This generated a distribution of group averaged Az values based on 2000 randomised data sets, from which we could estimate the Az value leading to a significance level of p<0.01.

## Results

### Psychometric behaviour and perceptual thresholds

Subjects’ performance was quantified by fitting behavioural performance (“first stream higher” responses) with psychometric curves to derive psychometric thresholds (σ) and points of subjective equality (PSE, μ).

Figure 2A shows the group-level psychometric curves for each sensory condition. On unisensory trials, thresholds were significantly lower (i.e. better performance) for high compared to low reliability stimuli across subjects (p<0.05, Table I). Thresholds were comparable for the auditory and both congruent audio-visual conditions (p>0.05, Table I). Thresholds on audio-visual trials were however significantly lower compared to the visual conditions (p<0.05, Table I). This demonstrates that performance was comparable on audio-visual and auditory trials, and better for high vs. low reliable stimuli.

**Figure 2.**
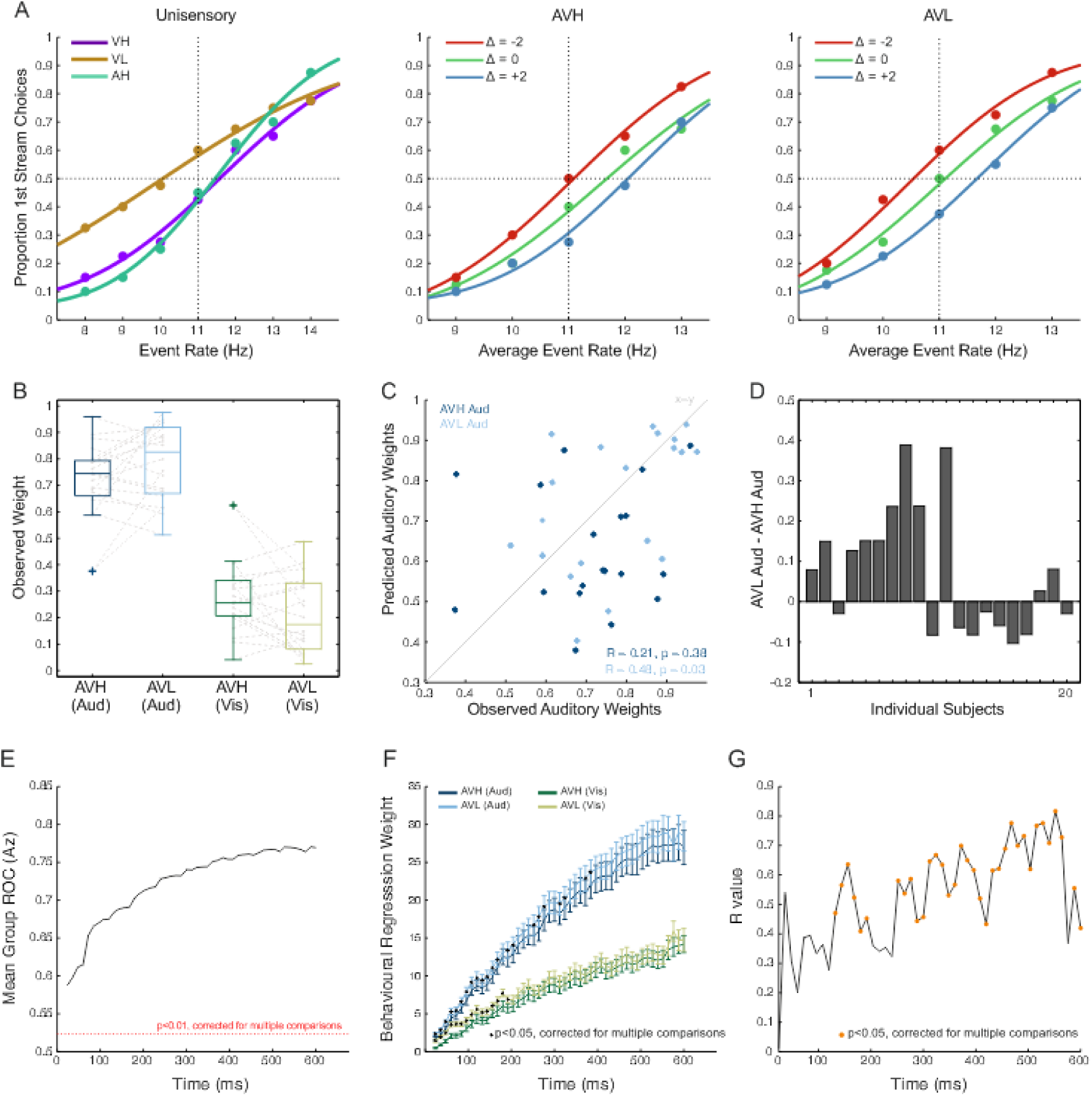
Behavioural Results. (A) Group (n=20) level psychometric curves are displayed as the proportion of “first stream” decisions as a function of event rate for each condition. Note that for incongruent trials the x-axis indicates the average event rate (Δ= Visual Rate – Auditory rate). Vertical dashed lines represent the standard rate (11 Hz) and horizontal dashed lines represent chance (50%) performance. (B) Observed perceptual weights with individual subject data shown in grey. AVH represents the audiovisual condition where both the auditory and visual cues were equally reliable. AVL represents the audiovisual condition where the auditory was highly reliable and the visual was less reliable. (C) Predicted and observed auditory weights separately for AVH and AVL conditions. Subjects with data below the grey line (representing WPRED = WOBS) have higher auditory weights than predicted. (D) The difference between auditory weights in the AVH and AVL conditions (WAVL – WAVH) for each subject. (E-F) Logistic regression was used to predict single trial choice (event rate >/<11 Hz) based on the accumulated event rate at each time point in the trial. (E) Performance of the logistic model quantified using the area under the ROC (dashed line p<0.01, randomisation test) (F) Auditory and visual perceptual weights derived from the regression model.Time points with significant reliability effects are denoted with black circles. (G) Correlation of perceptual weights derived from psychometric curves and from the logistic model. Time points with significant correlations are marked with orange circles.

**TABLE I.** Analysis of psychometric data.

| Psychometric Fits |  |  | Comparison of Perceptual Thresholds |  |  |  |
| --- | --- | --- | --- | --- | --- | --- |
| | Median $\sigma$ | Median $\mu$ | | Z value | p value | Effect Size |
| AUD | 2.19 | 11.45 | VH vs AUD | -2.688 | 0.007 | -0.425 |
| VH | 2.87 | 11.44 | VL vs. AUD | -3.658 | 0.0003 | -0.579 |
| VL | 4.55 | 10.53 | VH vs. VL | -3.136 | 0.002 | -0.496 |
| AVH ($\Delta=0$) | 1.98 | 11.72 | AUD vs. AVH | -0.336 | 0.737 | -0.053 |
| AVH ($\Delta=+2$) | 1.80 | 12.25 | AUD vs. AVL | -0.261 | 0.794 | -0.041 |
| AVH ($\Delta=-2$) | 1.89 | 11.05 | AVH vs. AVL | -0.018 | 0.986 | -0.003 |
| AVL ($\Delta=0$) | 2.29 | 11.14 | AVH vs. VH | 3.322 | 0.0009 | 0.525 |
| AVL ($\Delta=+2$) | 1.82 | 11.94 | AVH vs. VL | 3.621 | 0.0003 | 0.573 |
| AVL ($\Delta=-2$) | 2.10 | 10.58 | AVL vs. VH | 3.397 | 0.0007 | 0.535 |
|  |  |  | AVL vs. VL | 3.919 | 0.00009 | 0.619 |

Comparing psychometric curves for congruent and incongruent multisensory conditions showed that regardless of visual reliability, subjects preferentially weighted the auditory modality, as demonstrated by shifts in the psychometric curves towards the auditory rate (Fig. 2A *left and right).* However as expected, this shift was more pronounced in the low reliability condition, showing a stronger influence of the auditory modality when visual reliability was reduced (Table I).

Based on these psychometric data we derived a set of predicted and observed perceptual weights for each modality under each level of reliability. Predicted auditory weights significantly increased when visual reliability was reduced (p<0.05, Table II). However, this pattern was not consistently found in the observed weights (Fig. 2B), where there was no significant difference between observed auditory weights between reliabilities (p>0.05, Table II). Furthermore, a direct comparison between observed and predicted weights (Fig. 2C) revealed only a weak correlation (AVH p>0.05, AVL p<0.05, Table II).

**TABLE II.** Analysis of predicted and observed psychometric perceptual weights.

| High vs. Low Reliability Perceptual (Auditory) Weights |  |  |  | Predicted vs. Observed Perceptual Auditory Weights |  |  |
| --- | --- | --- | --- | --- | --- | --- |
| | Z | p-value | Effect Size | | $R_s$ | p-value |
| Pred AVH vs. Pred AVL | -2.837 | 0.005 | -0.454 | Pred AVH vs. Obs AVH | 0.21 | 0.381 |
| Obs AVH vs. Obs AVL | -1.261 | 0.207 | -0.199 | Pred AVL vs. Obs AVL | 0.48 | 0.034 |

The lack of a significant difference between unisensory and multisensory thresholds and the weak correlation between observed and predicted weights suggests that observers did not systematically follow the behavioural pattern predicted by Bayesian models of multisensory integration. This is corroborated by Figure 2D, which shows the magnitude and direction of the weight shift across subjects. Only 11 subjects showed a shift of auditory weight shifts in the predicted direction (i.e. increased auditory weighting when visual reliability is reduced). For the present study, this heterogeneity in the change of perceptual weights with reliability presents a unique opportunity to investigate the neural correlates of perceptual weights independently of an effect of sensory reliability, as these two effects are dissociable across subjects.

### Evolution of Perceptual Weights over Time

To obtain insights into the temporal dynamics of the perceptual weighting process we modelled the relationship between behavioural choice and sensory evidence at each time point within a trial, and derived a set of dynamic perceptual weights for each modality and reliability condition (*see Methods*). Importantly, having a time-resolved measure of perceptual weights allowed us to test when during a trial these changed with sensory reliability.

We found that our measure of sensory evidence (accumulated rate) was significantly predictive of behavioural choice across the trial (randomisation test, p<0.01, Fig. 2E). Auditory and visual perceptual weights increased as sensory evidence was accumulated throughout the trial, and confirmed that subjects preferentially weighted the auditory over the visual modality (cluster randomisation test, p<0.05, Fig. 2F, Table III). We found that both auditory and visual weights changed significantly with reliability (cluster randomisation tests, p<0.05, Fig. 2F), and did so early during the trial (five auditory clusters covering 24ms to 384ms, one visual cluster covering 24ms to 192ms, Table III).

**TABLE III.**
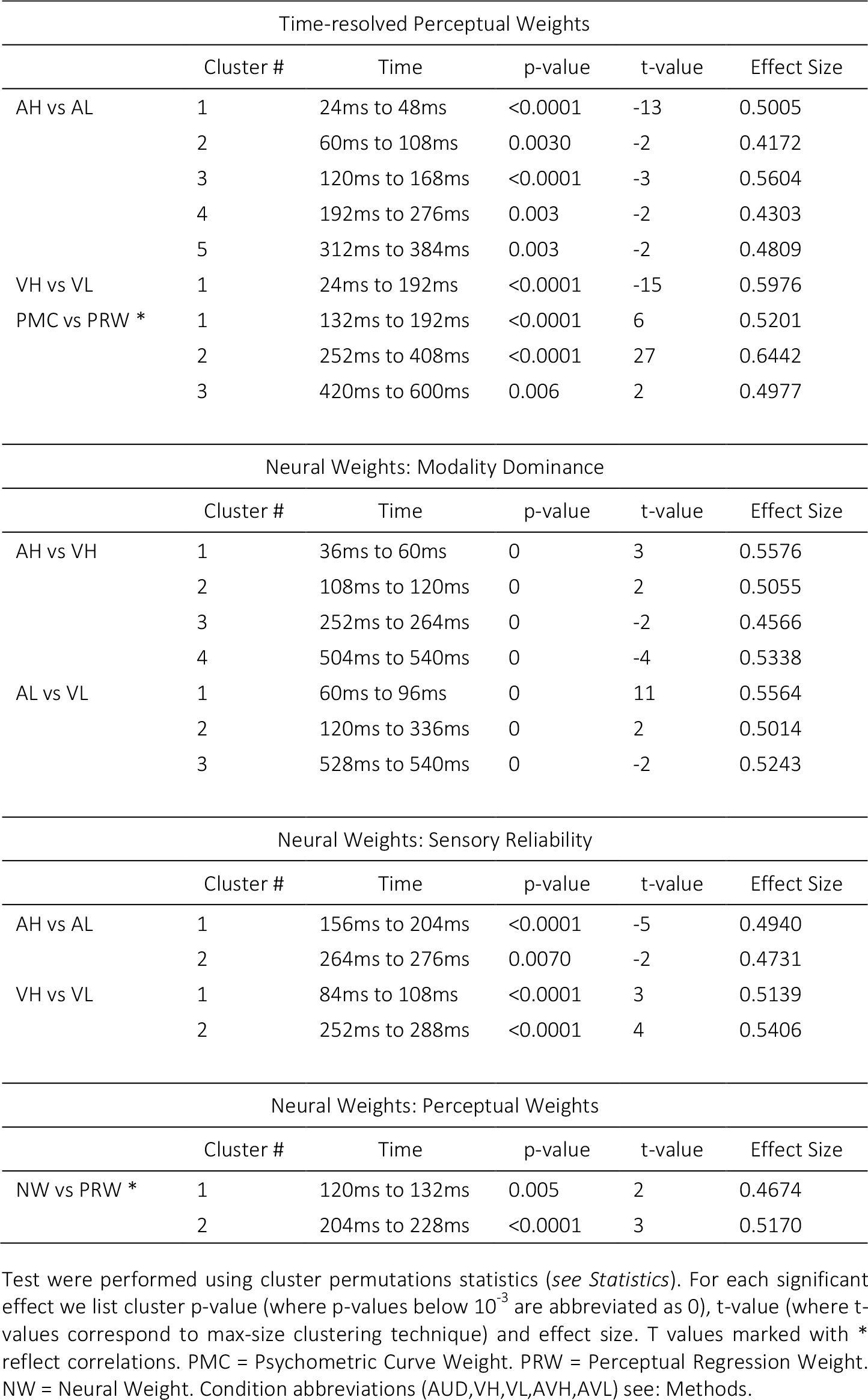
Statistical effects for comparisons of perceptual or neural weights between conditions

To test how consistent these time-resolved perceptual weights were with those derived from the psychometric curves, we computed their correlation. Significant correlations (cluster randomisation tests, p<0.05, Fig. 2G) emerged during three epochs that collectively covered most of the trial (three clusters: 132ms to 192ms, 252ms to 408ms, and 420ms to 600ms, Table III).

### EEG signatures of event rates

We used a linear discriminant analysis to extract EEG components that maximally discriminated between event rates (</> 11 Hz). Such an approach allowed the use of the discriminant output (Y) as a proxy to the single trial stimulus evidence reflected in the EEG activity [Kayser et al., 2016; Philiastides et al., 2014; Ratcliff et al., 2009], and we exploited this to link the neural signature of the sensory input to changes in the external sensory reliability and perceptual weights (*see* Methods).

Figure 3A displays the discriminant performance across subjects. Significant performance emerged early in the trial (48ms to 396ms, randomisation test, p<0.01) and was highest during three time epochs (96ms to 120ms, 168ms to 204ms, and 252ms to 288ms, with peaks defined as Az performance >0.58). To assess how this neural signature of the event rate was modulated by sensory reliability, we regressed the discriminant output (Y) against the accumulated rate to obtain neural sensory weights.

**Figure 3.**
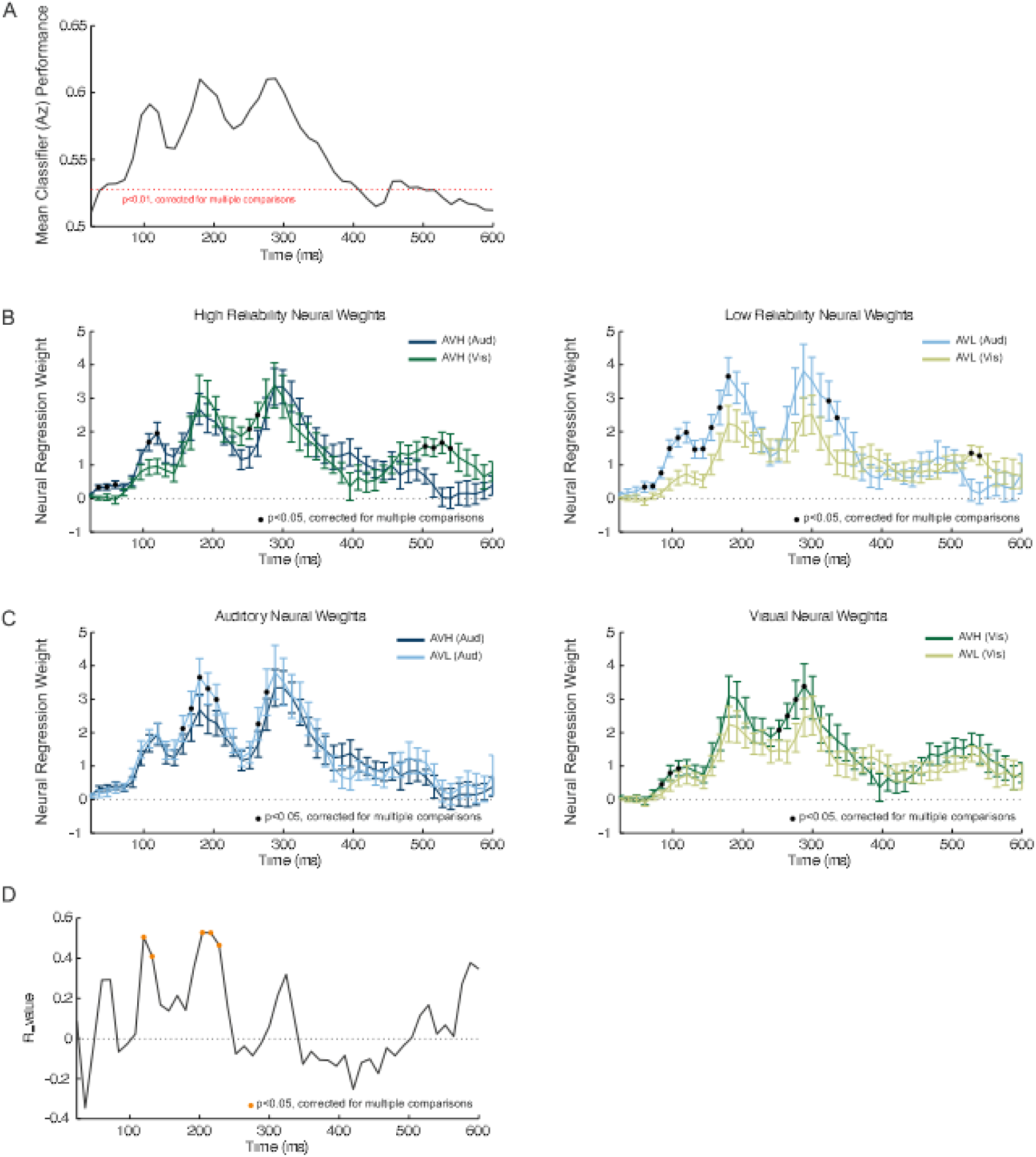
EEG Decoding, Neural Weights and Neuro-Behavioural Correlation. (A) Group averaged performance of a linear classifier discriminating between the two stimulus conditions (event rate >/<11Hz), quantified using the area under the ROC curve. The discriminant output (Y) was calculated using a sliding time window of 55ms aligned to the window onset, from 24ms to 600ms post stimulus onset. (B) Neural weights for each modality for high reliability (left) and low reliability trials (right). (C) Neural weights for each reliability for auditory (left) and visual stimuli (right). In each panel (B,C), time points with significant reliability / modality effects are indicated by black circles. (D) Neuro-behavioural correlation between the perceptual and neural weights obtained from the regression models. Time points with significant correlations are indicated by orange circles.

### Influence of reliability on neural weights

First, to determine whether the neural weights exhibited a similar bias towards the auditory modality as the perceptual weights, we compared them between modalities. Indeed, auditory weights dominated in both reliability conditions (Fig. 3B *left* and *right*), and these difference emerged at several epochs across the trial (four clusters AVH: 36ms to 60ms, 108ms to 120ms, 252ms to 264ms, and 504ms to 540ms; three clusters AVL: 60ms to 96ms, 120ms to 336ms, 528ms to 540ms; cluster randomisation tests, p<0.05,Table III).

Second, we quantified how neural weights were affected by sensory reliability. Consistent with perceptual weights, auditory weights were significantly higher when the visual reliability was reduced, and these differences emerged during two epochs (156ms to 204ms, and 264 to 276ms; cluster randomisation test, p<0.05;

Fig. 3C *left;* Table III). In addition, visual weights were significantly lower when the visual reliability was reduced at two epochs (84ms to 108ms, and 252ms to 288ms; cluster randomisation test, p<0.05, Fig. 3C *right;* Table III).

Finally, we asked whether there was a significant relationship between the reliability effect on the time-resolved perceptual and the neural weights. This revealed two epochs during which neural and perceptual weights correlated significantly: 120ms to 132ms, and 204ms to 228ms (cluster randomisation test, p<0.05, Fig. 3D; Table III).

Summarising the above results in order of time (rather than by statistical contrast) shows that there is an evolving pattern of weights as the trial progresses. Starting from stimulus onset, we first observe a change in visual weights (starting 84ms) and a significant relationship between perceptual and neural weights (starting 120ms). This is followed by a change in auditory weights (starting 156ms) and another epoch where there is a significant relationship between perceptual and neural weights (starting 204ms). Finally, we observe a change in both visual (starting 252ms) and auditory weights (starting 264ms), later in the trial.

These three statistical contrasts above revealed six epochs during which neural weights exhibited patterns of interest in relation to our main question. To disentangle whether these epochs represented distinct neural processes, or whether some of these likely relate to the same underlying neural generators, we analysed the relationship between these effects further. We did so by comparing scalp projections and neural weights between the six epochs (Supplementary Fig. 1). This revealed that temporally adjacent topographies (84ms to 108ms and 120ms to 132ms; 158ms to 204ms and 204ms to 228ms; and 252ms to 288ms and 264ms to 276ms) were highly correlated (within Epochs: R_S_ >0.6, p<0.005 for all comparisons). The reliability difference in neural weights (Eqn. 3) at temporally adjacent epochs were also highly correlated (R_s_ >0.6, p<0.001) and showed similar patterns of neural weights. Therefore, we report the topographies and neural weights for these three time epochs of interest (Epoch 1: 84ms to 132ms; Epoch 2: 158ms to 228ms; and Epoch 3: 252ms to 276ms).

Figure 4 shows the neural weights and forward model scalp topographies for the three epochs of interest. The first component was characterized by a scalp projection revealing strong contributions of occipital electrodes, consistent with a potential origin in sensory cortices. The second component had a scalp projection that revealed strong contributions from fronto-central electrodes, consistent with sensory (temporal) and parietal regions. Finally, the third component revealed strong contributions from temporal and central electrodes.

**Figure 4.**
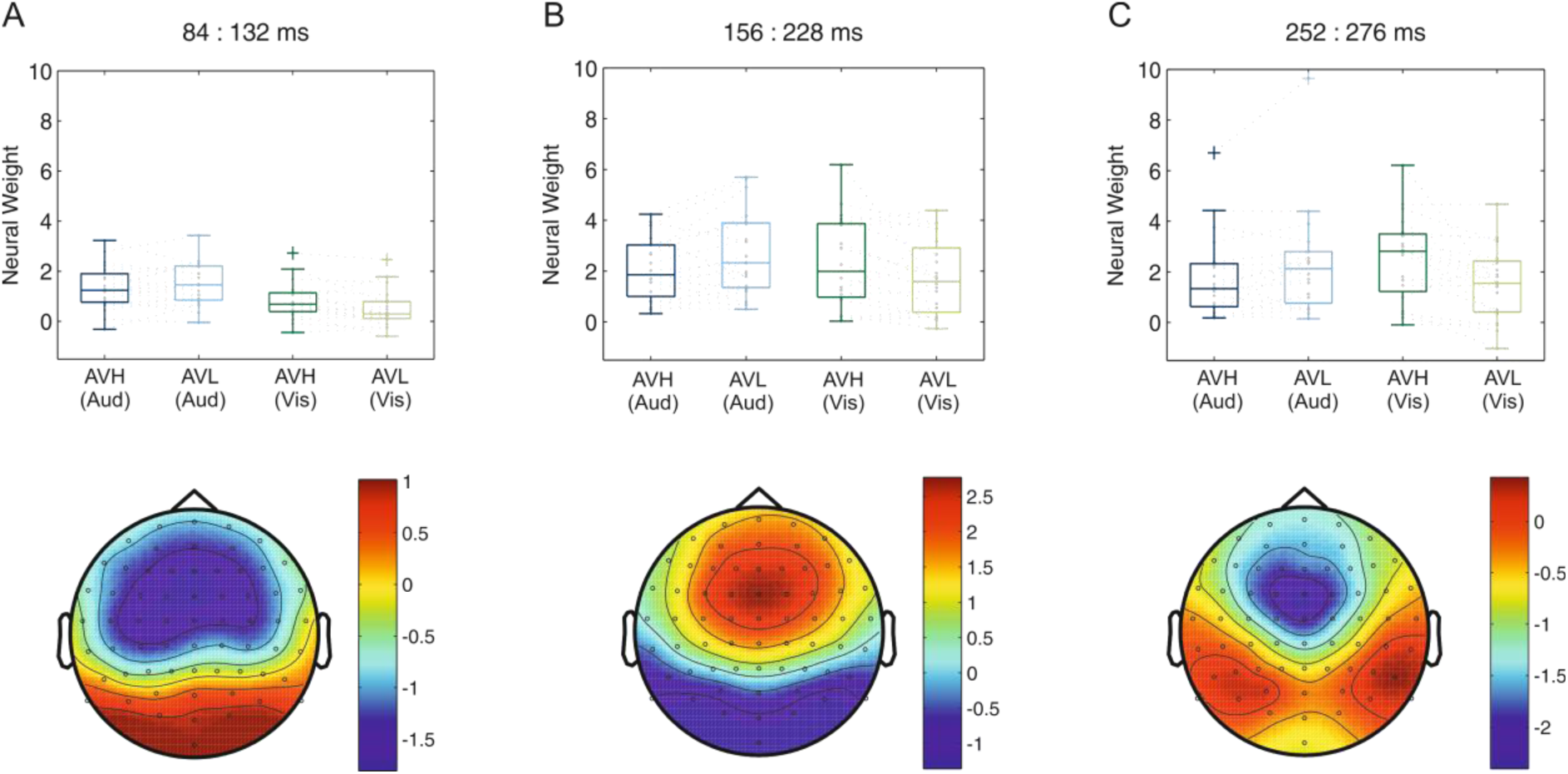
Neural Weights and Scalp Topographies for three EEG components of interest. Each component was defined based on the statistical contrast between sensory reliabilities (Fig. 3), or a significant neuro-behavioural (N2B) correlation (Fig. 3D). In each panel, boxplots represent neural weights averaged over each epoch, with individual subject data in grey. Topographies represent the group averaged forward models averaged over the epoch, where the values represent the correlation between the discriminating output (Y) and the underlying EEG activity.

## Discussion

This study examined the temporal dynamics of reliability based cue weighting during an audio-visual task. Our results revealed three epochs during the integration process in which brain activity exhibited correlates of sensory reliability and perceptual weights. Specifically, perceptual and neural weights are modulated by the reliability of sensory information as early as 84ms after stimulus onset and correlate with the perceptual weights underlying perceptual choice around 120ms. Together, these results suggest that the weighted combination of sensory information occurs early and within sensory regions, rather than only late and in amodal association cortices. Additionally, we step beyond previous neurophysiological [Fetsch et al., 2012; Gu et al., 2008; Morgan et al., 2008] and neuroimaging studies [Beauchamp et al., 2010; Helbig et al., 2012; Rohe and Noppeney, 2015b; Rohe and Noppeney, 2016] by revealing the temporal evolution of the sensory weighting process in functionally specific brain activity. Finally, by dissociating the influence of sensory reliability on the representation of sensory information from perceptual weighting in EEG responses rather than demonstrating a simple modulation of evoked response amplitudes, we show that these early effects reflect sensory and computationally specific processes.

### Early Effects of Sensory Reliability

We found that early during the trial neural sensory weights scaled with reliability. At the earliest window (84ms to 132ms) these effects were associated with changes in visual weights, while at the slightly later window (starting at 156ms) auditory neural weights scaled with changes in visual reliability. At a slightly later time window (starting at 252ms) changes in auditory and visual weights were evident. This finding that visual and auditory weights first scaled with reliability at different latencies during the trial is noteworthy. While visual weights were affected early (<100ms), auditory weights increased with decreasing reliability of the visual stimulus later (around 150ms). This temporal dissociation of visual and auditory weight changes with reliability could reflect the adaptive nature of multisensory integration during this paradigm. Perhaps visual encoding is adjusted at short latencies and in a bottom-up (i.e. sensory driven manner) to cope with trial-by-trial changes in visual sensory reliability; in contrast, auditory encoding is adjusted only later (possibly as result of top-down processes) in order to meet the increased demands for representing the unreliable sensory environment.

### Early Correlates of Perceptual Weighting

In our behavioural data, we show that the time-varying perceptual weights are significantly predictive of choice, and show similar reliability differences as seen in perceptual weights obtained from the psychometric curves. These differences again emerge early in the trial, starting around 24ms after stimulus onset. In addition, we show that neural correlates of the perceptual weighting process emerge early in the trial (around 120ms after stimulus onset), and a long time before the perceptual choice at the end of the trial.

We were able to dissociate these processes, given that sensory reliability and perceptual weights were not strongly associated across participants, as not every subject attributed perceptual weights in a statistically optimal manner. Hence, the scaling of sensory representations in proportion to the physical reliability of the sensory input, and the correlation of neural with perceptual weights are computationally distinct and so reflect different aspects of the sensory-perceptual cascade. It remains unclear whether these perceptual weights are adjusted on each trial individually, and in response to the experienced sensory reliabilities, or whether they are at least in part already established based on task-context in a predictive manner even before stimulus onset. Future work is required to elucidate the precise neural correlates of these perceptual weights and how different brain regions contribute to establishing the perceptual integration process.

### Temporal organisation of effects

The temporal organization and localisation of the reliability and perceptual weighting effects in three clusters showed distinct patterns of topographies. At the earliest window (84ms to 132ms), effects of sensory reliability and perceptual weights were associated with scalp topographies consistent with early sensory areas. At the slightly later time point (starting at 156ms), our effects were associated with activity over central electrodes consistent with activations including prominent contributions from auditory cortex. Finally, at the latest window (252ms to 288ms), effects were associated with activity over posterior and central regions. While the localization of the relevant EEG components was quite distributed, our results fit with the notion that earliest effects arise from occipital sensory regions and are followed by activity in the temporal and parietal lobe.

This evolving pattern of activation lends support to the idea that early sensory and parietal regions encode unisensory cues and represent the integrated evidence weighted by the relative reliability and scaled by task-demands and relevance. This complements existing findings from fMRI work showing multisensory interactions occurring along primary sensory and parietal areas in response to changing reliability [Helbig et al., 2012; Beauchamp et al., 2010; Rohe & Noppeny 2015, 2016]. Taken together with this prior literature, our results support the idea that sensory reweighting is an evolving and hierarchical process, with multisensory interactions emerging along the sensory pathway in primary sensory and parietal areas. Yet our results add a temporal dimension to these processes and demonstrate that effects related to external sensory reliability and perceptual weighting emerge at slightly different times and from distinct brain regions.

## Acknowledgements

We would like to thank Joachim Gross for advice on the analysis and Marios Philiastides for help with implementing the decoding algorithms. This work was supported by the European Research Council (to C.K. ERC-2014-CoG; grant No 646657) and a BBSRC DTP studentship (to S.C.B.).

## Supplementary Figures and Tables

**Supplementary Figure 1.**
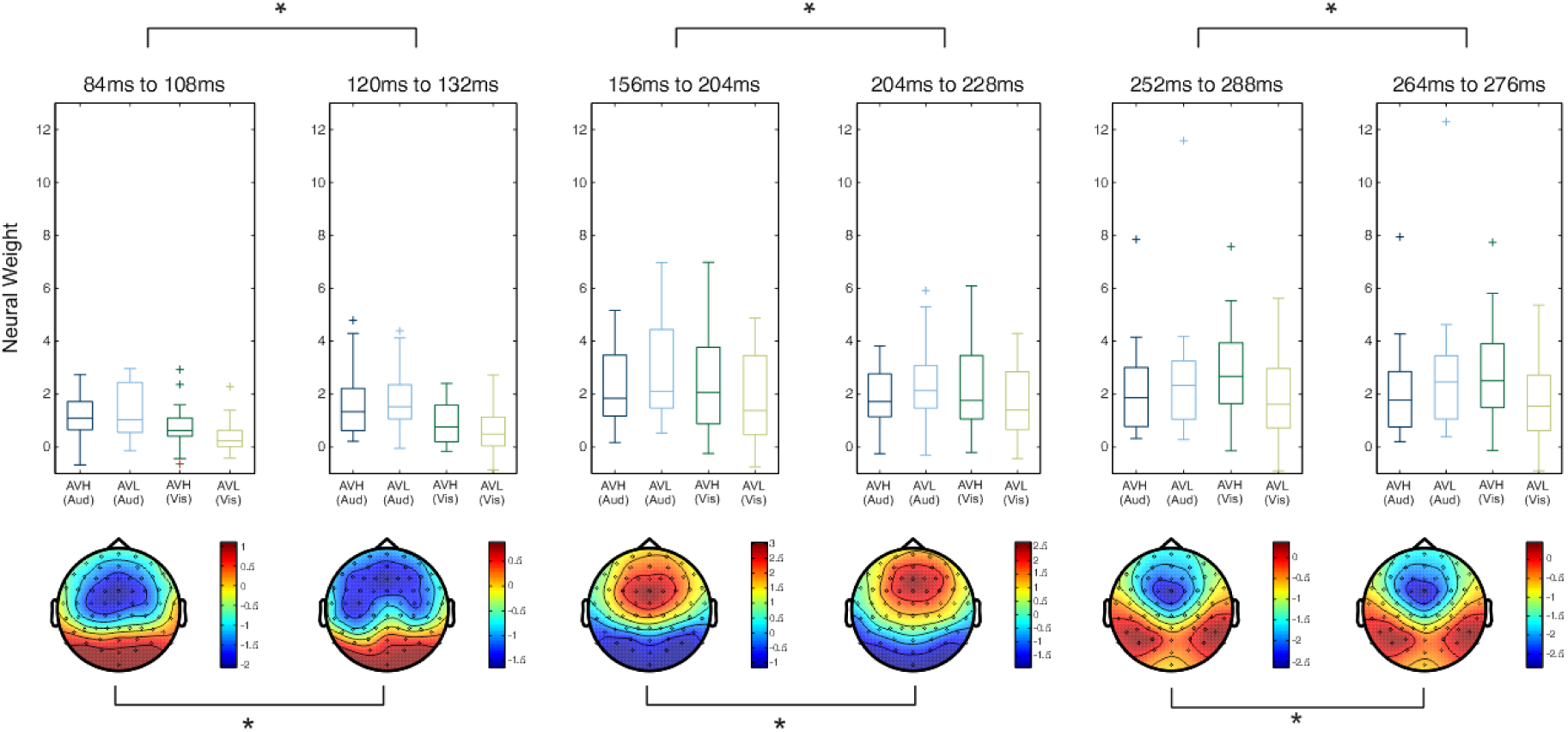
Neural weights and scalp topographies underlying the six time epochs that showed a significant effect of reliability. Topographies marked with * represent R >0.6 and p<0.005 for correlations between scalp topographies. For each epoch the difference in neural weights (AVHAUD-AVHVIS)-(AVLAUD-AVLVIS) was calculated and correlated, * represent R>0.6 and p<0.001.

